# The liver clock tunes transcriptional rhythms in skeletal muscle to regulate mitochondrial function

**DOI:** 10.1101/2025.01.17.633623

**Authors:** Valentina Sica, Tomoki Sato, Ioannis Tsialtas, Sophia Hernandez, Siwei Chen, Pierre Baldi, Pura Muñoz Cánoves, Paolo Sassone-Corsi, Kevin B. Koronowski, Jacob G. Smith

## Abstract

Circadian clocks present throughout the brain and body coordinate diverse physiological processes to support daily homeostasis and respond to changing environmental conditions. The local dependencies within the mammalian clock network are not well defined. We previously demonstrated that the skeletal muscle clock controls transcript oscillations of genes involved in fatty acid metabolism in the liver, yet whether the liver clock also regulates the muscle was unknown. Here, we use hepatocyte-specific *Bmal1* KO mice (Bmal1^hep-/-^) and reveal that approximately one third of transcriptional rhythms in skeletal muscle are regulated by the liver clock vivo. Treatment of myotubes with serum harvested from *Bmal1*^hep-/-^ mice inhibited expression of genes involved in metabolic pathways, including oxidative phosphorylation. Overall, the transcriptional changes induced by liver clock-driven endocrine-communication revealed from our in vitro system were small in magnitude, leading us to surmise that the liver clock acts to fine-tune metabolic gene expression in muscle. Strikingly, treatment of myotubes with serum from *Bmal1*^hep-/-^ mice inhibited mitochondrial ATP production compared to WT and this effect was only observed with serum harvested during the active phase. Overall, our results reveal communication between the liver clock and skeletal muscle–uncovering a bidirectional endocrine communication pathway dependent on clocks in these two key metabolic tissues. Targeting liver and muscle circadian clocks may represent a potential avenue for exploration for diseases associated with dysregulation of metabolism in these tissues.

## INTRODUCTION

Circadian rhythms are adaptive mechanisms that temporally regulate our physiology to anticipate environmental changes associated with the 24-hour light-dark cycle. At the cellular level, circadian rhythms are driven by biological clocks that consist of a series of interlocked transcriptional-translational feedback loops that regulate rhythmic gene expression of clock-controlled genes. In mammals, the transcription factor BMAL1 is a primary non-redundant regulator of this system (Bunger et al., 2000), and mouse models using specific depletion or reconstitution of *Bmal1* have played key roles in revealing a core function of clocks in regulation of metabolism.

Though local roles are increasingly well defined, how clocks coordinate metabolism systemically remains poorly understood. This warrants further investigation since perturbation of circadian rhythms is increasingly implicated as a contributory factor to diverse diseases including metabolic syndrome and cancer (Deota and Panda, 2020; Masri and Sassone-Corsi, 2018).

The liver, a hub of systemic metabolism, displays extensive circadian regulation. We recently used a mouse model of *Bmal1* reconstitution in the liver of otherwise clockless mice (Greco et al., 2021; Koronowski et al., 2019) and found that whilst the cooperation of the liver clock with feeding-fasting rhythms is essential to maintain carbohydrate homeostasis, redox and lipid metabolism requires the additional contribution of the skeletal muscle clock (Greco et al., 2021). From simultaneous reconstitution of *Bmal1* in liver and muscle, we then showed that communication between clocks in these two tissues in the presence of feeding fasting rhythms is the minimal clock network required to ensure systemic glucose tolerance (Smith et al., 2023). We previously identified effects of the muscle clock on metabolic gene expression in the liver, yet it remained unclear how the liver clock influences the skeletal muscle.

Here, we investigated the specific role of the liver clock on skeletal muscle rhythmicity by using a mouse model of *Bmal1* deletion in hepatocytes. Overall, we found 30% of the 24h-rhythmic transcriptome in muscle is regulated by the liver clock, and that serum-dependent signalling plays a defining role in this process. Assessment of metabolic function in myotubes then revealed a functional impact of this signalling axis on mitochondrial respiration. Overall, this study adds further evidence supporting the role of peripheral clocks in inter-organ crosstalk.

## METHODS

### Animals

Mice were bred and housed at the University of California, Irvine (UCI) vivarium, in accordance with the guidelines of the Institutional Animal Care and Use Committee (IACUC) at UCI. Animal experiments were designed and conducted with consideration of the ARRIVE guidelines whenever possible. Male 8 to 14-week-old mice were group housed under a standard 12 hour light–12 hour dark light cycle, fed ad libitum with vivarium chow, and randomly assigned to groups. Mice were entrained for 2 weeks prior to tissue collection at different diurnal timepoints. Bmal1 hepatocyte-specific knockout mice (*Bmal1*^hep-/-^) were generated by crossing Bmal1-flox mice^115^ with Alfp-Cre. Experimental genotypes were: 1. wild type (WT) – *Bmal1^wt/wt^*, *Alfp-*cre^tg/0^; and 2. *Bmal1*^hep-/-^– *Bmal1^flox/flox^*, *Alfp-*cre^tg/0^. Male 6 to 8 week-old mice for providing serum for seahorse experiments were bred and housed at the University of Texas Health San Antonio animal facility. Mice were fed a standard chow diet ad libitum (7912 Teklad Irradiated LM-485 Mouse/Rat Diet) and were group housed in standard home cages within circadian-controlled cabinets (Actimetrics). Environmental conditions included 23.9°C ± 1.5°C temperature, humidity monitoring at approximately 30% and the light intensity at the cage level is measured at approximately 100 lux using a luminometer. Lighting schedules were fully automated using the programmable lighting features of the cabinets, ensuring minimal disturbance to the animals except for routine tasks, such as bedding changes and the replacement of food and water. All light schedules consisted of daily periods of 12 hr of light and 12 hr of dark.

### Transcriptomics

Muscle:Liquid nitrogen-frozen gastrocnemius muscle was crushed into a homogeneous powder and RNA was extracted using DirectZol (Zymo) columns. Myotubes: 24 hours after treatment, myotubes were washed 2x with PBS on ice. RNA was then extracted on harvested myotubes using TRIzol and DirectZol (Zymo) columns. Extracted RNA was sequenced by Novogene (Novogene Co) for total RNA. Quality checking was performed using FastQC v0.11.8 and to mm10 (STAR--outFilterMultimapNmax 20 --alignSJoverhangMin 8 –alignSJDBoverhangMin1 -- outFilterMismatchNmax 999 --outFilterMismatchNoverReadLmax0.04 -- alignIntronMin 20--alignIntronMax 1000000 --alignMates-GapMax 1000000 -- outSAMattributes NH HI NM MD). RNA-seq quality assessment was performed using RSeQC v3.0.1 tool.

### Circadian and differential gene expression analyses

DryR (Weger et al., 2021) (https://rdrr.io/github/naef-lab/dryR/) was run in drylm mode on FPKM data; BICW>0.6 and amplitude >0.25 were used as minimum requirements for inclusion of a gene in a given model. LimoRhyde (Singer and Hughey, 2019) was used to differential regulation of 24h rhythmic genes identified with JTK_CYCLE (Hughes et al., 2010); p<0.01 was used to identify rhythmic genes, as in our previous study (Greco et al., 2021). The BioVenn (Hulsen et al., 2008) web tool (https://biovenn.nl/) was used to perform venn-diagram overlaps between genes identified in the DryR and LimoRhyde analyses. For differential gene expression analysis in myotubes, limma (Ritchie et al., 2015) was used on FPKM data. Genes with zero values for any sample were excluded. P<0.01 was used as a cut off to identify significant genes. Volcano plots were generated using ggplot2 (Wickham, 2009). ChatGPT 4o Canvas Mode (accessed January 2025) was used to generate and edit scripts to use Limma (Ritchie et al., 2015), Bioconductor (Huber et al., 2015) and ggplot2 (Wickham, 2009) inside R (R 4.4.1 GUI 1.80 Big Sur Intel build 8416). Galaxy (The Galaxy Community, 2024) was used to join tables using the web interface usegalaxy.org.

### Myotube cultures and serum treatment

For sequencing experiments performed at UCI, C2C12 mouse skeletal myoblasts (ATCC; CRL-1772) were grown in Dulbecco’s modified Eagle’s medium (DMEM, 11965-092) (4.5 g/l D-glucose and Glutamate, pyruvate free) (GIBCO) and 15 % fetal bovine serum (FBS; GIBCO) with 1% penicillin–streptomycin. At 90% confluence, C2C12s were differentiated for myotubes using DMEM and 2% horse serum (GIBCO) with 1% penicillin-streptomycin. Media change on 30% of the media was performed each day, and were differentiated for 4 days. Mouse serum to be used as treatment for the myotubes was removed from -80 storage and thawed on ice. Serum for individual or pooled for 2 mice was made up to 25% v/v in Dulbecco’s modified Eagle’s medium (DMEM, 11965-092) (4.5 g/l D-glucose and Glutamate, pyruvate free) (GIBCO) with 1% penicillin–streptomycin. After filter sterilisation, media/serum mixes were added (each well receiving biologically distinct serum), cells returned to the incubator, then harvested 24 hours later. For Seahorse experiments performed at UTHSA, C2C12 cells were plated in high glucose (4.5 g/L) DMEM, supplemented with 10% FBS and 1% P/S, at a density of 40.000 cells/cm2, in a XFe96/XF Pro 96-well cell culture microplate. At 90 – 100% cell confluency, ∼1.5 to 2 days after plating, C2C12 myoblasts were washed twice with prewarmed PBS and incubated with high glucose (4.5 g/L) DMEM, supplemented with 2% horse serum and 1% P/S (differentiation media) for 4 days. The media was replaced every 2 days, retaining 10% of the previous media to maintain C2C12-secreted factors essential for growth and differentiation. The fourth day after the change in differentiation media, C2C12 myotubes were treated for 24hrs with 1% serum derived from WT and Bmal1hep-/- ZT20 mice. We opted to incubate with 1% v/v serum (rather than 25% v/v) to match more closely the serum percentage in the culture media.

### Seahorse analyses on myotubes

The Agilent Seahorse XF Cell Mito Stress Test was performed on C2C12 cells above, according to the manufacturer’s instructions. After the completion of the assay, protein measurement in each well of the 96 well-plate was performed using the BCA assay and was utilized for the quantification of the results. Of note, protein measurement and subsequent normalisation was not conducted in the first of the three biological replicates.

### Western blot

Western blot was performed from whole-cell lysates as in (Koronowski et al 2019 Cell). Briefly, liver was homogenized in RIPA lysis buffer (50 mM tris-HCl [pH 8], 150 mM NaCl, 5 mM EDTA, 15 mM MgCl2 and 1% NP-40) supplemented with a cocktail or protease and HDAC inhibitors. Samples were let on ice for 30 min and then sonicated at 40% amplitude, 4 cycles of 5 sec on, 5 sec off. Samples were centrifuged for 15 min at max speed and protein content of the supernatant was determined by Bradford assay. Protein was separated by SDS-PAGE on 8% gels, transferred to a nitrocellulose membrane, and blocked with 5% milk TBST (0.1% Tween-20 in TBS) for 2 hours at RT. Primary antibodies were diluted in blocking buffer and incubated with membranes overnight at 4°C (BMAL1, Abcam – ab93806; Tubulin). Membranes were incubated with HRP-conjugated secondary antibodies in blocking buffer for 1 hour at RT, developed with HRP substrate, and visualized via autoradiography film.

### qPCR on liver samples

To extract RNA, livers were homogenized in TRIzol. RNA, DNA, and protein layers were separated with chloroform and RNA was precipitated with a standard isopropanol-ethanol protocol. Purified RNA was resuspended in DNase- and RNase-free water and quantified with by nanodrop. The Maxima First Strand cDNA Synthesis Kit (Thermo) was used to retrotranscribe 500 ng of RNA into cDNA. Real-time quantitative polymerase chain reaction (qPCR) was performed with a QuantStudio 3 (Applied Biosystems) and a PowerUp SYBR Green (Applied Biosystems) reaction mixture. Expression was normalized to 18S control. Primer sequences were as follows: 18s For - 5′-CGCCGCTAGAGGTGAAATTC-3′; 18s Rev 5′-CGAACCTCCGACTTTCGTTCT-3’; Arntl (Bmal1) For - 5′-GCAGTGCCACTGACTACCAAGA-3′; Arntl Rev - 5′-TTGCAATCTTACCCCAGACA-3′.

### Statistical analyses

Unless otherwise indicated, all data are presented as individual datapoints (each data point refers to a biological replicate). Information regarding sample size, statistical test, and significance can be found in the corresponding figure legend or in the results text. Methods sections describe statistical analyses of large-scale datasets. Field standards were applied to determine sample sizes for circadian RNA-seq experiments. Prism 6.0 (GraphPad) was used to graph/illustrate data and perform statistical analyses.

### Data availability

FPKM lists are provided as supplemental files (muscle data-Table S1, C2C12 data-Table S4), alongside DryR (Table S2), LimoRhyde (Table S3) and limma (Tables S5-S7) output data.

## RESULTS

### Core clock rhythms in skeletal muscle are independent from the liver clock

To assess the role of the liver clock in cross-tissue signalling to skeletal muscle, we generated hepatocyte-specific *Bmal1* KO mice (Bmal1^FL/FL^; AlfPCre^+/-^; referred to as Bmal1^hep-/-^) and WT controls (Bmal1^wt/wt^; AlfPCre^+/-^). We first confirmed loss of *Bmal1* at both the transcript and protein level in livers from Bmal1^hep-/-^ mice (Figure S1A-B). We then performed RNA sequencing on gastrocnemius muscle harvested from adult male WT and Bmal1^hep-/-^ mice at 4 hour timepoints spanning the 12h light–12h dark cycle (zietgeber time ZT0 (start of light phase), ZT4, ZT8, ZT12, ZT16, ZT20; n=3 mice per timepoint per genotype) (Figure 1A; Table S1). Assessment of expression profiles of core clock genes in skeletal muscle did not reveal obvious impairment of the core clock transcriptional loop (Figure 1B, Figure S1C). This apparent insensitivity of the muscle clock to deletion of *Bmal1* in the liver was further supported via the use of the *dryR* algorithm to classify rhythmic genes, which returned all core clock genes in the same model, relating to maintained rhythmicity in both genotypes (Figure 1C). We next used the JTK_CYCLE algorithm as a further approach, and found comparable p values for oscillation of each gene in Bmal1^hep-/-^ and WT mice (Figure 1D). Peak phase and amplitude of detected oscillations were reported as unaltered in *dryR* (Figure 1C), and were no changes were detected by JTK_CYCLE (Figure 1E and 1F). We therefore summarise that the oscillation of the core clock transcriptional loop in skeletal muscle is independent from the liver clock.

**Figure 1:**
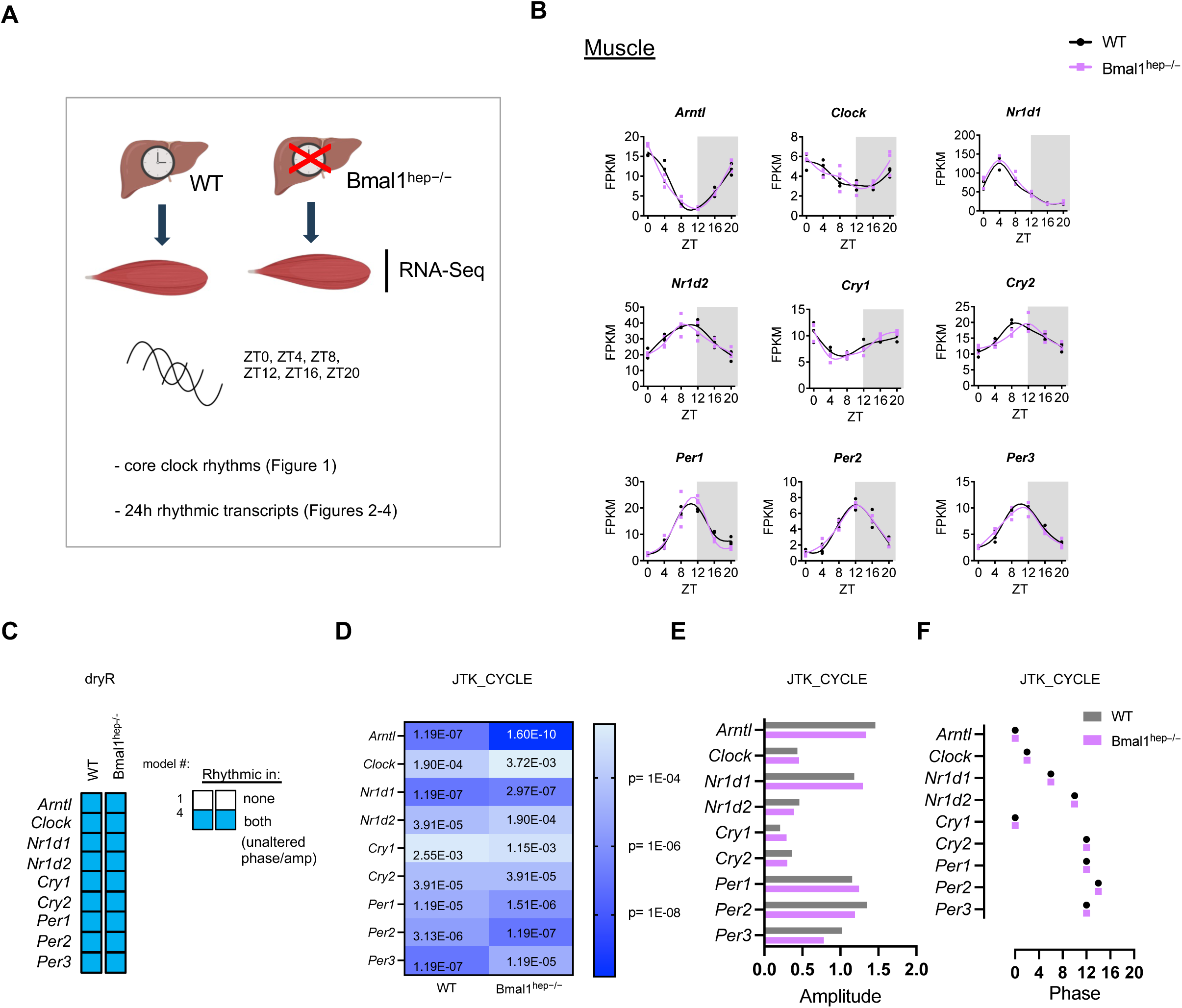
Core clock rhythms in skeletal muscle are independent from the liver clock. A. Schematic of experimental design. ZT; zeitgeber time. B. Expression of core clock genes in skeletal (gastrocnemius) muscle isolated from WT and hepatocyte specific *Bmal1* KO (Bmal1^hep-/-^) mice (n=3 male mice per genotype per timepoint), FPKM; Fragments Per Kilobase of transcript per Million mapped reads. C. Rhythmicity of core clock genes in skeletal muscle by DryR analysis. D-F. Rhythmicity (D), peak phase (E) and amplitude (F) of core clock genes in skeletal muscle by JTK_CYCLE analysis.

### The hepatocyte clock controls a subset of transcriptional rhythms in skeletal muscle

In order to reveal the extent of genome-wide impacts of the liver clock on transcriptional rhythms in gastrocnemius muscle, we used *dryR* to classify 24h oscillatory genes into different models based upon their rhythmicity in each genotype (Figure 2A; Table S2). Using this approach, we found that 69.5% (1295/1863) of oscillatory genes in skeletal muscle are refractive to deletion of the liver clock (Figure 2A-C). Nevertheless, we detected 14.7% of genes lose (273/1863) and 14.1% gain (262/1863) oscillation in the muscles of Bmal1^hep-/-^ versus WT mice, in addition to 1.7% (33/1863) of genes that oscillate in both genotypes but with different amplitude and/or phase (Figure 2B-C; Figure S2A). Thus, approximately one third (30.5%) of the muscle diurnal transcriptome is regulated by the liver clock. Whilst genes resilient to liver clock deletion display a bimodal distribution of peak phases around ∼ZT0 and ∼ZT12, genes that lose oscillation are characterised by more broadly distributed peak phases (Figure 2C) and a higher overall amplitude (Figure S2A). Genes that gain oscillation mostly peak in the dark phase around ZT20, and exhibit a lower amplitude versus resilient or lost *dryR* models (Figure 2C, Figure S2A).

**Figure 2:**
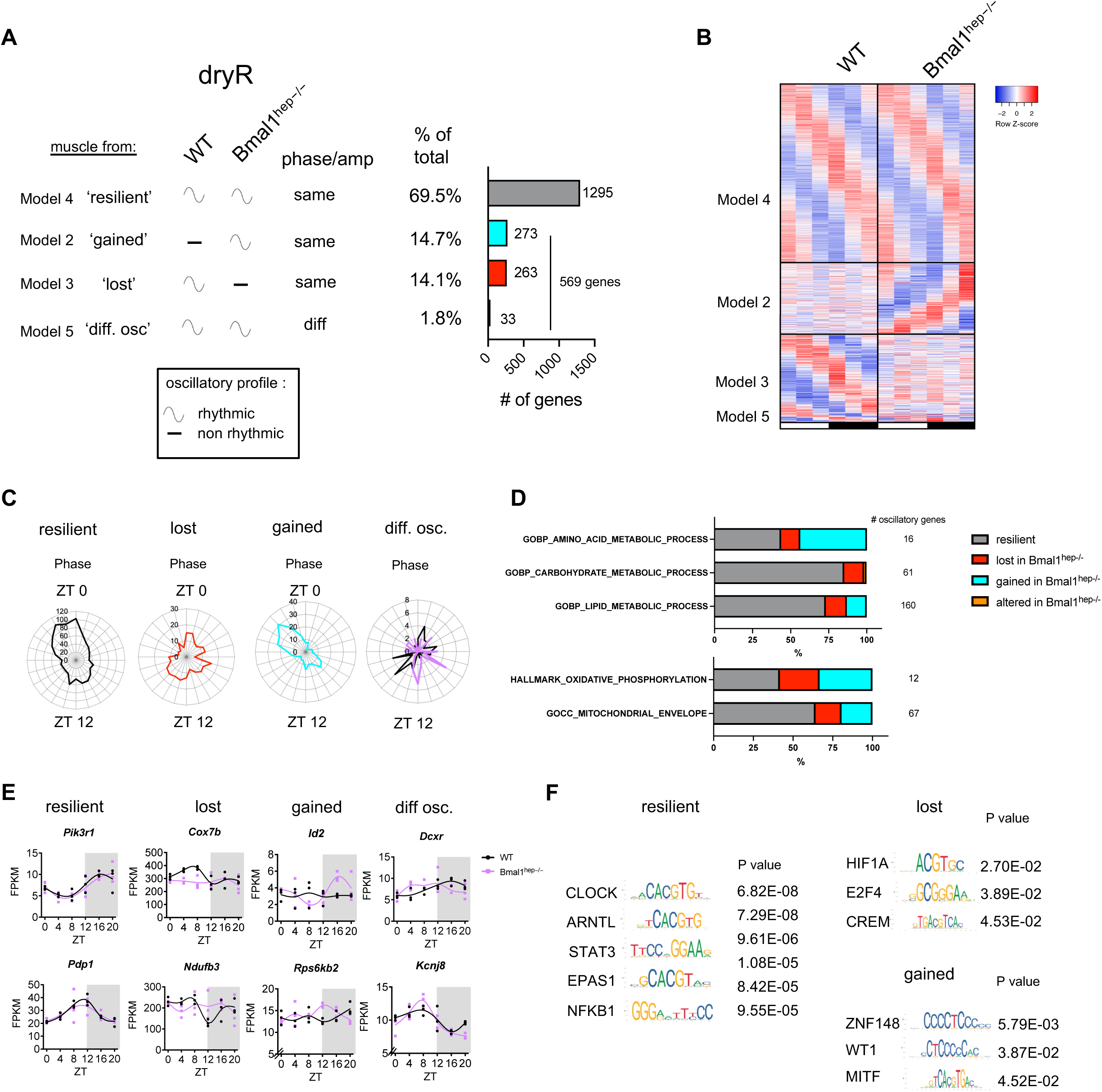
The hepatocyte clock controls a subset of transcriptional rhythms in skeletal muscle. A. dryR analysis on gastrocnemius muscle harvested from hepatocyte specific Bmal1-KO mice (Bmal1^hep-/-^; see also Figure 1A). B. Phase-aligned heatmaps of genes in specified dryR classes (BICW >0.6, amp<0.25). C. Phase plots for each category D. Gene ontology analysis E. Example genes F. candidate TFs (TRRUST Transcription Factors 2019). Motif images from https://jaspar.elixir.no/.

Seeking to define the characteristics of genes that are resilient versus regulated by the liver clock, we performed KEGG pathway enrichment analysis for each gene class (Figure S2B). Genes resilient to influence from the liver clock (1295) were enriched for circadian rhythm as well as terms related to insulin signalling (KEGG terms: Insulin resistance, AGE-RAGE signalling in diabetic complications, MAPK signalling pathway) amongst others (Figure S2B). We also did not observe significant changes in gene expression of Slc2a4 (encoding for GLUT4, the main glucose transporter in skeletal muscle involved in insulin-induced glucose uptake) and Myod1 (Figure S2C), which has previously been identified as a factor involved in circadian gene expression in muscle (Hodge et al., 2019). For de novo oscillating genes (273), KEGG pathway enrichment scores were much weaker and no significant enrichments were detected for genes that either lose oscillation in Bmal1^hep-/-^ mice (262) or exhibited differential oscillations (33). Hence, the effect of liver clock deletion appears to be broad rather than targeted to particular pathways.

To gain insight into the effect on key metabolic pathways with relevance for skeletal muscle, we classified the rhythmic behaviour of genes involved in amino acid, carbohydrate and lipid metabolism, alongside mitochondrial function–a feature of skeletal muscle previously shown to be under regulation of the muscle clock (Kumar et al., 2024) (Figure 2D-E). Across these pathways, between 14.8% to 58.3% of oscillatory genes in skeletal muscle were regulated by the liver clock (Figure 2D). Oscillatory genes involved in carbohydrate metabolism were most resilient to liver clock deletion (85.2% captured in model 4). For the 26.9% (43/160) of lipid metabolic oscillatory genes, approximately half lost oscillations in the muscle in the absence of the liver clock (22/43) and half gained de novo oscillations (21/43). Though few oscillatory genes (16) were identified as involved in amino acid metabolism, 56.3% (9/43) were affected upon liver clock knockout–7 losing oscillation and 2 exhibiting de novo rhythmicity. Interestingly, we also found that for terms involved in mitochondria metabolism such as GOCC (Gene Ontology Cellular Component) Mitochondrial Envelope (also encompasses all genes in the GOCC Mitochondrial Membrane Category) and Oxidative Phosphorylation, 24/67 (35.8%) and 7/12 (58.3%) were affected, respectively, and genes were almost evenly distributed between the lost and gained classifications. Several cytochrome c oxidase (*Cox*) and NADH dehydrogenase (*Nduf*) complex genes, which constitute the electron transport chain, lost oscillation in Bmal1^hep-/-^ muscle (Figure 2E). In line with the lack of detected effect on the core clock (Figure 1), CLOCK and ARNTL were the top transcription factors predicted to be involved in regulation of resilient genes (Figure 2F). STAT3 and NFκB, key transcription factors involved in regulation of skeletal muscle mass, were also indicated as candidate transcription factors in this class. Less significant p values were observed for the lost and gained class of genes (none for the differentially regulated gene class). HIF1α, whose interaction with the clock in skeletal muscle regulates mitochondrial and glycolytic metabolism (Peek et al., 2017), was the top predicted transcription factor for genes that lose oscillation in muscles of Bmal1^hep-/-^ mice. ZNF148, a transcription factor downregulated during muscle development (Passantino et al., 1998) was predicted to regulate de novo oscillating genes in the muscle of Bmal1^hep-/-^ mice.

We also performed analysis with a differential circadian rhythmicity analysis tool, LimoRhyde (Figure S3; Table S3) which identified fewer rhythmic genes in skeletal muscle overall (352) yet 84.4% (297/352) were also identified in the *dryR* analysis (Figure S2A-B). Overall, 68/352 (19.3%) of oscillatory genes in muscle were identified as regulated by the liver clock by this approach (Figure S3B). The loss of high amplitude rhythms in the muscle of Bmal1^hep-/-^ mice that was observed in our *dryR* analysis (Figure S2A) was also found via this approach, as well as the gain of de novo low amplitude rhythms (Figure S2C). LimoRhyde identified genes with a loss of oscillation peak around ZT14 whilst de novo genes that peak at ZT3 (Figure S3D). In agreement with our dryR data, several metabolic genes and mitochondrial genes were still captured as differentially-regulated using this approach (Figure S3E-F). Use of the same tool and thresholds also allowed comparison to our previous study that assessed communication from the skeletal muscle clock to liver (Greco et al., 2021). LimoRhyde identified 2316 oscillatory genes in liver (Greco et al., 2021) versus only 352 in skeletal muscle (Figure S3A), in agreement with our previous studies and other that revealed a higher rhythmicity in liver versus skeletal muscle (Smith et al., 2023).

### Serum signalling mediated by the hepatocyte clock regulates muscle gene expression

Liver and muscle are endocrine organs that exchange factors via the bloodstream in the form of proteins (hepatokines and myokines) and metabolites. Gene expression changes in muscle of Bmal1^hep-/-^ mice could be the result of changes in active circulating factors or other processes not mediated through the blood stream. To understand whether the liver clock uses the bloodstream to communicate with skeletal muscle, we used an in vitro setup in which we assessed gene expression via RNA-seq in C2C12 myotubes treated for 24 hours with serum from WT versus Bmal1^hep-/-^ mice collected either during daytime at ZT4 or nighttime at ZT16 (Figure 3A; Table S3). First, we compared the transcriptional response of myotubes to ZT4 or ZT16 serum from WT mice (Figure 3B, Table S5) and found 425 differentially expressed genes (DEGs). We note that the fold change of these changes tended to be relatively small (Figure S4), with DEGs expressing a mean FC of 1.16 (+/- 0.07 stdev) for the genes with higher expression at ZT4, and 1.27 (+/- 0.19 stdev) for genes with higher expression at ZT16. This is generally consistent with the effect of liver *Bmal1* knockout on the muscle diurnal transcriptome and fits with the notion that the liver tunes the muscle, rather than overtly alters it, under physiological conditions in healthy young-adult mice. We observed strong enrichments for ‘protein processing’ in genes with higher expression in myotubes treated with ZT4 serum (Figure 3C), and Ribosome in genes with higher expression in myotubes treated with ZT16 serum. We also observed significant enrichments for genes involved in metabolic pathways, N-glycan biosynthesis and fatty acid metabolism in genes with higher expression in myotubes treated with serum from ZT4. Oxidative phosphorylation stood out as the main metabolic pathway enriched in genes that exhibit a higher expression in myotubes treated with serum from ZT16.

**Figure 3:**
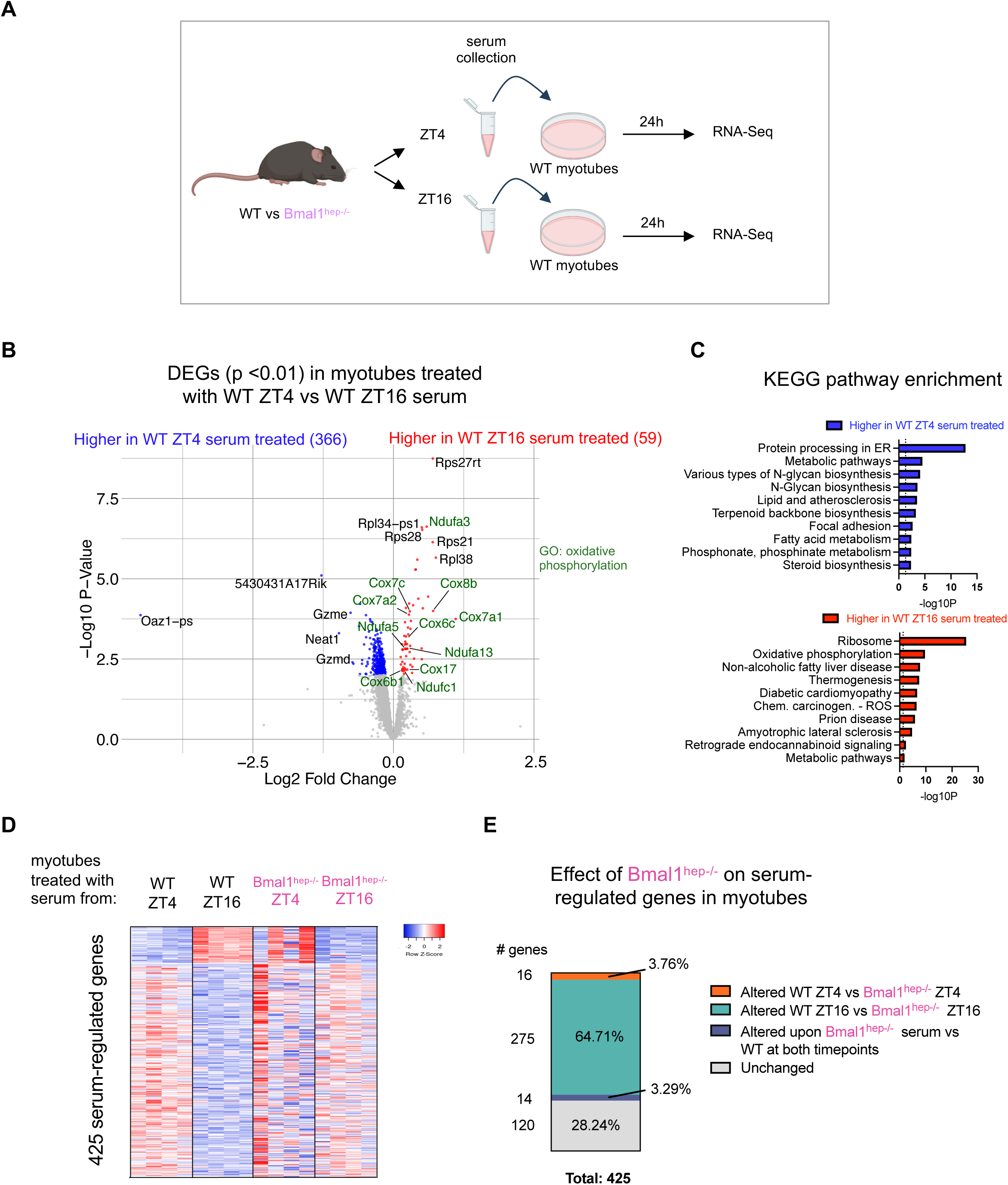
Serum signalling mediated by the hepatocyte clock regulates muscle gene expression. A. Experimental setup (see also Methods). Differentially expressed genes (DEGs) in C2C12-derived myotubes treated with ZT4 or ZT16 harvested serum from WT mice. C. KEGG pathway enrichment analysis for DEGs (p<0.01). Dotted line indicates p<0.05. D. Heatmap for serum-regulated genes showing expression in myotubes treated with serum from Bmal1^hep-/-^ mice. E. Effect of Bmal1^hep-/-^ on serum-regulated genes in myotubes. See also Figures S3-5.

Next, we probed this set of DEGs in myotubes treated with serum from Bmal1^hep-/-^ mice to assess the role of the liver clock in mediating these responses (Figure 3D). Strikingly, serum from Bmal1^hep-/-^ mice failed to replicate the ZT4 vs ZT16 DEGs observed in myotubes treated with WT serum. This was particularly evident in myotubes treated with ZT16 serum (Figure 3D). Indeed, directly comparing WT ZT16 serum-treated myotubes with Bmal1^hep-/-^ ZT16 serum-treated myotubes revealed that (64.71%) of the 425 WT serum DEGs were differentially expressed (Limma p<0.01, Figure 3E). Only 28.24% of the WT serum DEGs were statistically comparable between genotypes at either timepoint (Figure 3E; Tables S6-S7). These data indicate that the liver clock plays a role in tuning gene expression in skeletal muscle via serum-dependent factors.

### Hepatocyte clock-dependent endocrine signalling regulates mitochondrial respiration in myotubes

Gene expression changes that were reproducible in vivo and in vitro related to mitochondrial metabolism. In both experiments, we observed modulation of cytochrome c oxidase and NADH dehydrogenase complex genes, which are essential for electron transport chain and mitochondrial respiration. In vitro, we further expanded our analysis and assessed all Cox and Nduf gene expression in myotubes (Figure 4A). We found a consistent pattern, that these genes were upregulated in myotubes treated with ZT16 serum from WT mice but not Bmal1^hep-/-^ mice. We then asked whether these small but consistent changes would have physiological impacts on mitochondrial function. To that end, we measured mitochondrial respiration in serum-treated myotubes, using serum collected during nighttime (ZT20) to probe the functional aspects of the altered transcriptome that resulted from ZT16 serum treatment. Myotubes treated with serum from WT mice maintained higher rates of basal respiration (p=0.05) and ATP production (p<0.05) compared to myotubes treated with serum from Bmal1^hep-/-^ mice (Figure 4B). Non mitochondrial respiration was comparable between the groups (Figure 4B). Hence, our results indicate that the liver clock controls serum signalling during the dark phase to regulate mitochondrial function in muscle.

**Figure 4:**
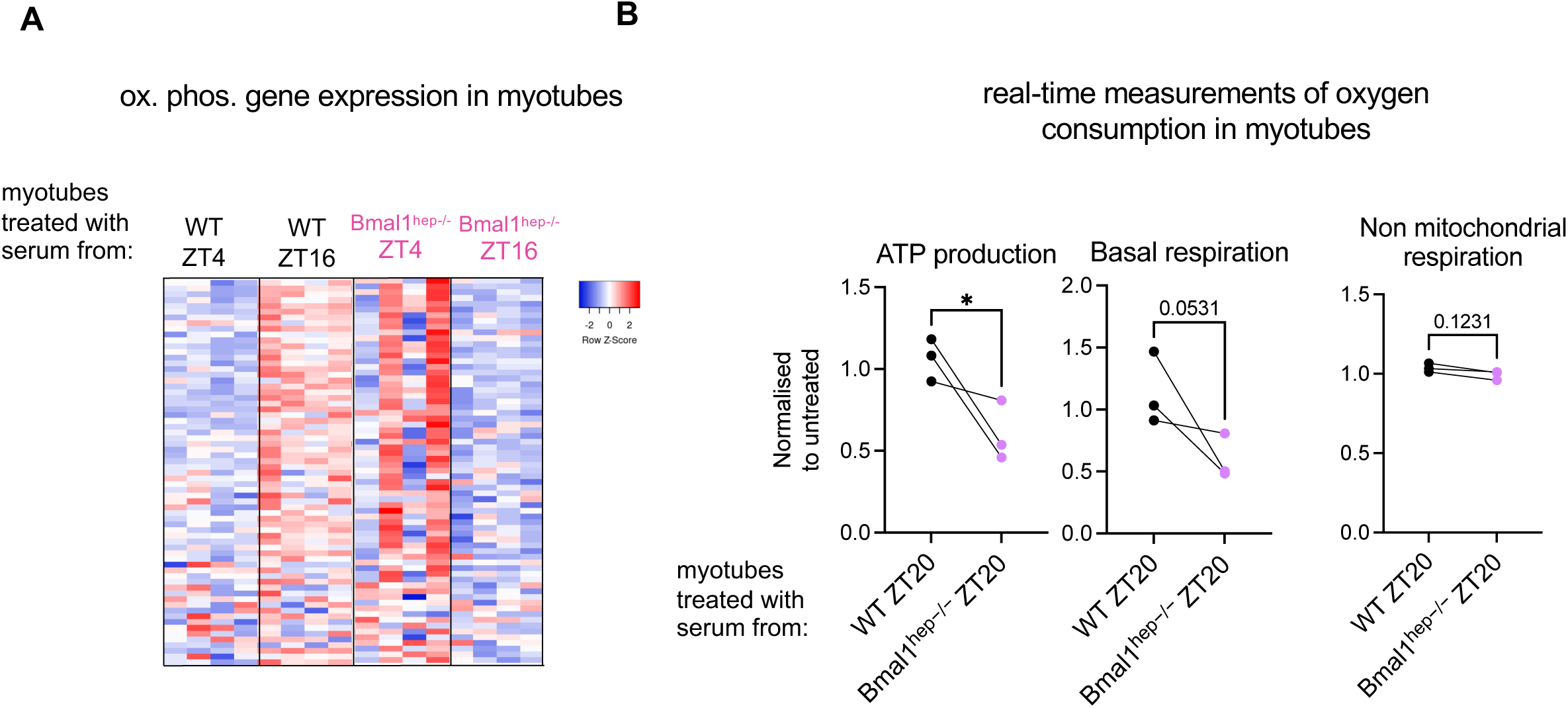
Hepatocyte clock-dependent endocrine signalling regulates mitochondrial respiration in myotubes. A. Heatmap of all Cox and Nduf genes detected in myotubes, with corresponding expression in each genotype and timepoint. B. Real-time measurement of oxygen consumption (Seahorse assay) showing ATP production in WT and Bmal1^hep-/-^-treated myotubes (n=3 biological replicates (serum samples), 1 biological replicate of each genotype assayed in each independent experiment). Each symbols refer to a biological replicate and lines indicate each independent experiment. *p<0.05 by Two-tailed unpaired t-test.

## DISCUSSION

Liver and muscle are critical metabolic tissues with key roles in glucose homeostasis and energy balance. In this study, we assessed the contribution of the intrinsic liver circadian clock on the circadian function in skeletal muscle. First, via analysis of the diurnal transcriptome of gastrocnemius muscle in WT versus Bmal1^hep-/-^ mice, we revealed that oscillations of core clock genes in muscle are not affected by the absence of the liver clock. This matches the lack of effect of the muscle clock on core clock oscillations in the liver that we observed in our previous studies (Greco et al., 2021; Smith et al., 2023). In addition, other groups have shown via deletion of clocks in either liver (Lamia et al., 2008; Manella et al., 2021), adipose tissue (Brooks et al., 2022) or gut (Chen et al., 2023) that there may be limited effects of peripheral clocks on the timing of clocks in other peripheral tissues.

By assessing overall 24h transcriptional rhythms in WT vs liver clock KO mice, we found that nearly one-third of oscillating genes in muscle were influenced by the liver clock. Based on comparison with our previous dataset (Greco et al., 2021), we estimate that this extent of regulation is similar to that observed in the opposite direction (by the muscle clock on liver). Hence, the cross-tissue circadian regulation by clocks in these two tissues in regard to magnitude of effect appears to be relatively balanced. Nevertheless, we observe differences in the type of pathways influenced by the liver clock on muscle and vice versa. Whereas genes involved in fatty acid metabolism are affected in livers from muscle clock KO mice (Greco et al., 2021; Viggars et al., 2024), muscle from liver clock KO mice instead show a more dispersed impact on metabolic gene expression.

In this study, we found that serum-derived signals under the control of the liver clock modulate mitochondrial gene expression and function in myotubes. This adds a new layer of complexity to our previous findings that the muscle and central clock cooperate to ensure mitochondrial health in muscle (Kumar et al., 2024). Whether these liver clock-dependent signals drive any diurnal differences in mitochondrial function in muscle is unknown.

In regard to the limitations of the current study, we note that this study is only using male mice, hence exploring sex-specific differences is likely of high relevance for future work. Furthermore, whilst our in vitro approach using treatment of myotubes with serum from WT and Bmal1^hep-/-^ mice revealed blood borne signalling was involved in communication between these tissues, the specific identity and nature of clock-dependent communicating factors remains to be determined.

Overall, we find that the dysregulation of a single peripheral tissue clock can affect metabolic regulation in a distal peripheral tissue, extending our knowledge regarding how peripheral clocks control metabolism. By defining a new route of inter-organ clock dependent signalling, this adds weight to a growing number of studies defining the impact of peripheral clocks in systemic communication. Investigating whether these clock-dependent signalling axes are dysregulated during disease may open new avenues to explore for therapeutic gain.

## Acknowledgments

V.S. acknowledges funding from FEBS long-term fellowship and European Union’s Horizon 2020 Research and Innovation Program under the Marie Sklodowska-Curie grant agreement 895380. The authors also acknowledge funding from MICINN-Spain (RTI2018-096068 to P.M.-C.), ERC-2016-AdG-741966 to P.M.-C., ‘‘la Caixa’’ (HEALTH-HR17-00040). UPGRADE (H2020825825), and María-de-Maeztu Program for Units of Excellence to UPF (MDM-2014-0370) and Altos Labs Inc. The work of S.C., M.S., and P.B. was in part supported by NIH grant GM123558. C.J. was supported by the AASLD Foundation Pinnacle Research Award in Liver Disease, the Edward Mallinckrodt, Jr. Foundation Award, and NIH/NIAAA R01 AA029124. The Koronowski lab is supported by the National Institute of General Medical Sciences of the National Institutes of Health under award number R35GM150618. Research in the J.G.S laboratory is supported by Agencia Estatal de Investigación (AEI) Aid RYC2022-035133-I funded by MICIU/AEI/10.13039/501100011033 and by the FSE+, and project PID2023-150233NA-100 funded by MICIU/AEI/10.13039/501100011033 and FEDER, EU, and AFM-Téléthon grant #28842. We thank Thomas Mortimer for providing a DryR script and advice on this analysis. Figures 1A and 3A use icons from BioRender.com.

---

**Figure S1:**
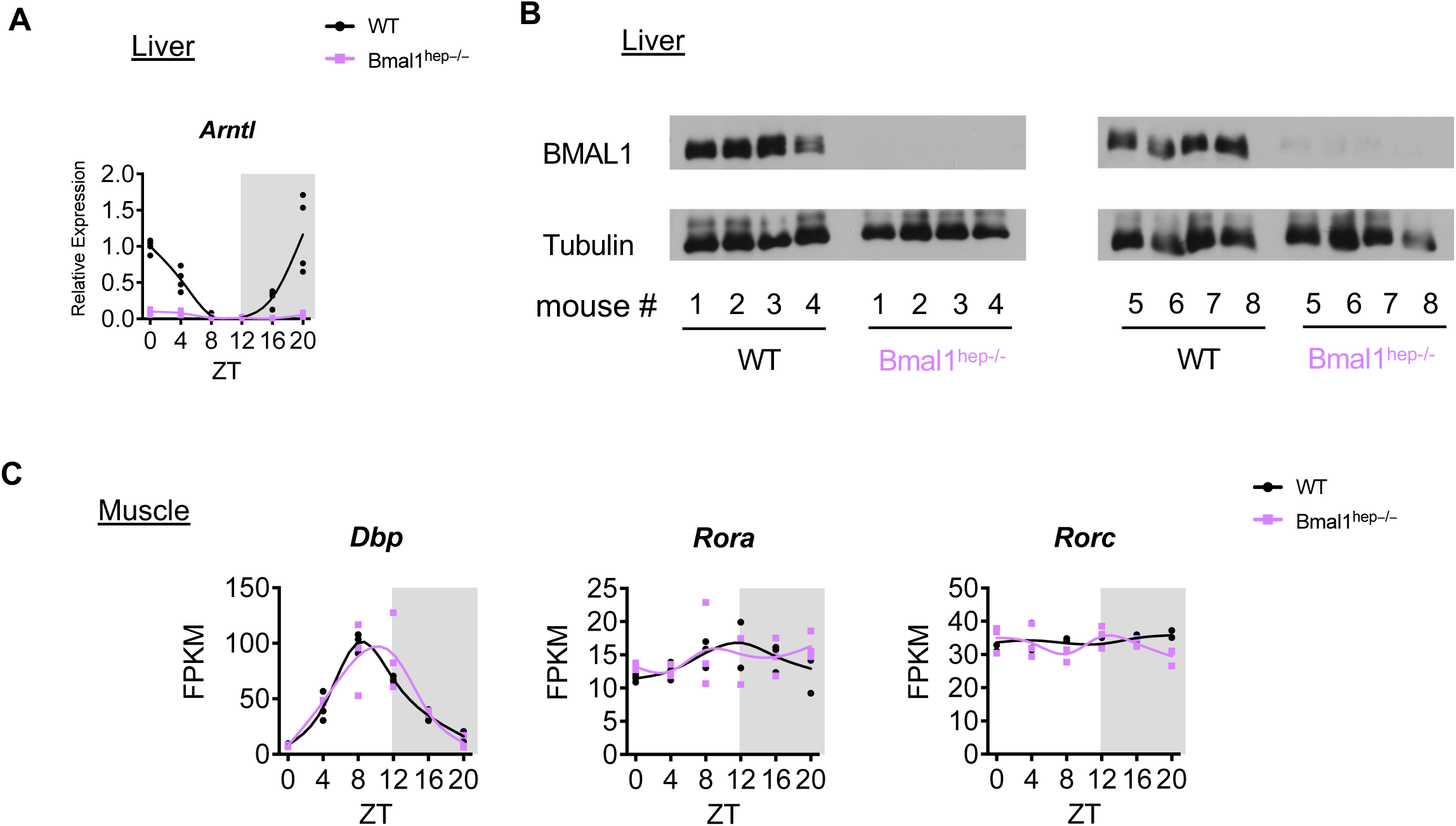
Supporting information for Figure 1 and 2. A. Confirmation of Arntl (Bmal1) deletion in livers of hepatocyte specific *Bmal1* KO (Bmal1^hep-/-^) mice (n=3 male mice per genotype per timepoint, qPCR). B. Western blot for BMAL1 in livers of hepatocyte specific *Bmal1* KO (Bmal1^hep-/-^) mice (n=8 mice per genotype). B. *Dbp* and *Rora/c* expression in muscle of WT and Bmal1^hep-/-^ mice. C. Expression of indicated genes in skeletal (gastrocnemius) muscle isolated from WT and hepatocyte specific *Bmal1* KO (Bmal1^hep-/-^) mice (n=3 male mice per genotype per timepoint), FPKM; Fragments Per Kilobase of transcript per Million mapped reads, ZT; zeitgeber time.

**Figure S2:**
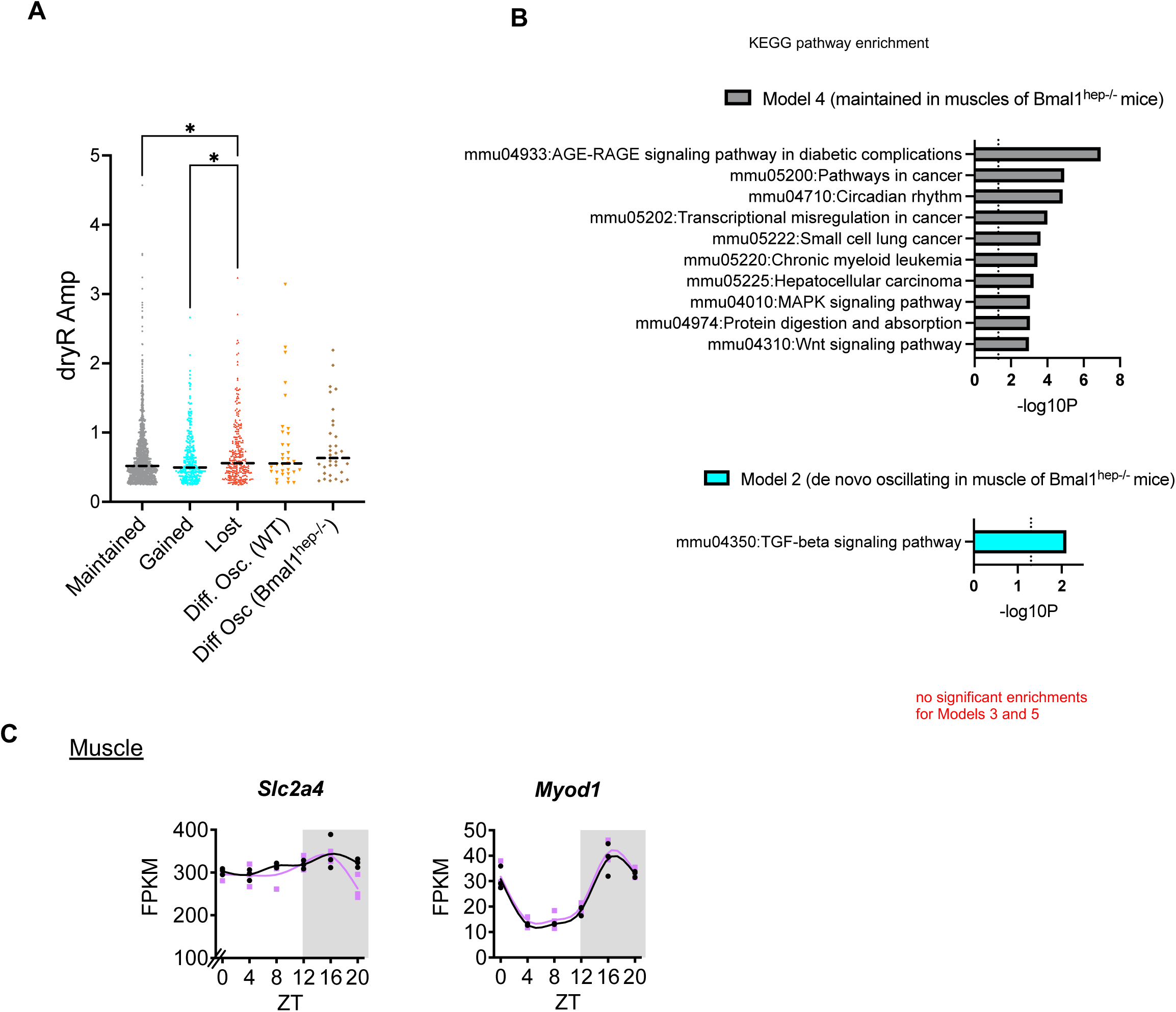
Supporting information for Figure 2. A) Amplitudes reported by DryR for each gene class) * p<0.05, ** p<0.01, Kruskal-Wallis test (Uncorrected Dunn’s-test). B) KEGG Pathway Enrichment Analysis (DAVID) for indicated gene classes. C) Expression of indicated genes in skeletal (gastrocnemius) muscle isolated from WT and hepatocyte specific *Bmal1* KO (Bmal1^hep-/-^) mice (n=3 male mice per genotype per timepoint), FPKM; Fragments Per Kilobase of transcript per Million mapped reads, ZT; zeitgeber time.

**Figure S3:**
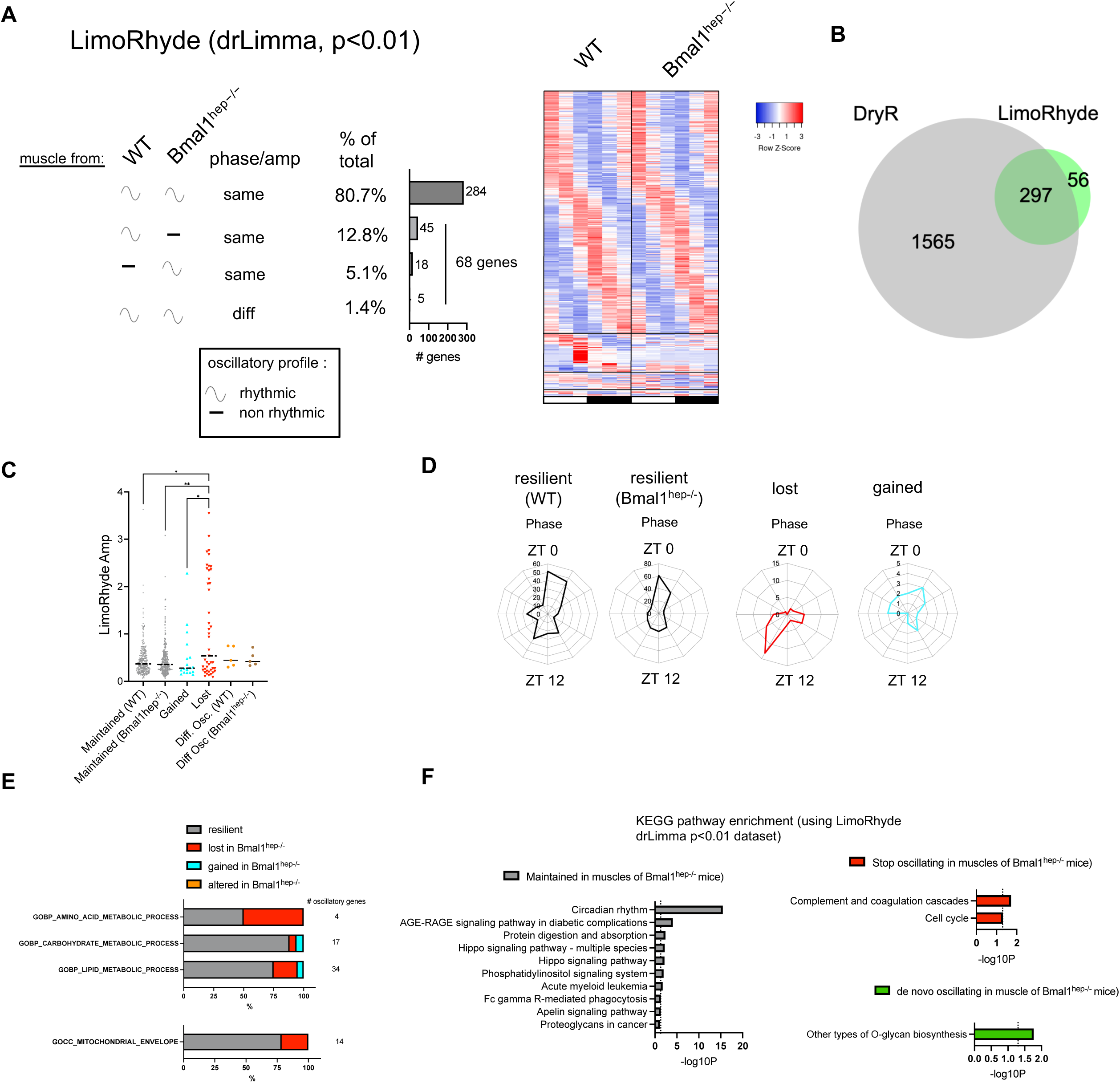
LimoRhyde rhythmicity analysis on muscles from from hepatocyte specific Bmal1-KO mice. A. LimoRhyde analysis using drLimma p<0.01 on transcriptome from gastrocnemius muscle harvested from hepatocyte specific Bmal1-KO mice (Bmal1^hep-/-^; see also Figure 1A). B. Comparison between dryR and LimoRhyde. C. Peak phase analysis. D. Amplitude analysis *p<0.05, ** p<0.01, Kruskal-Wallis test (Uncorrected Dunn’s-test). E. Gene ontology analysis F. KEGG pathway enrichment analysis for each gene class from drLimma p<0.01 LimoRhyde analysis.

**Figure S4.**
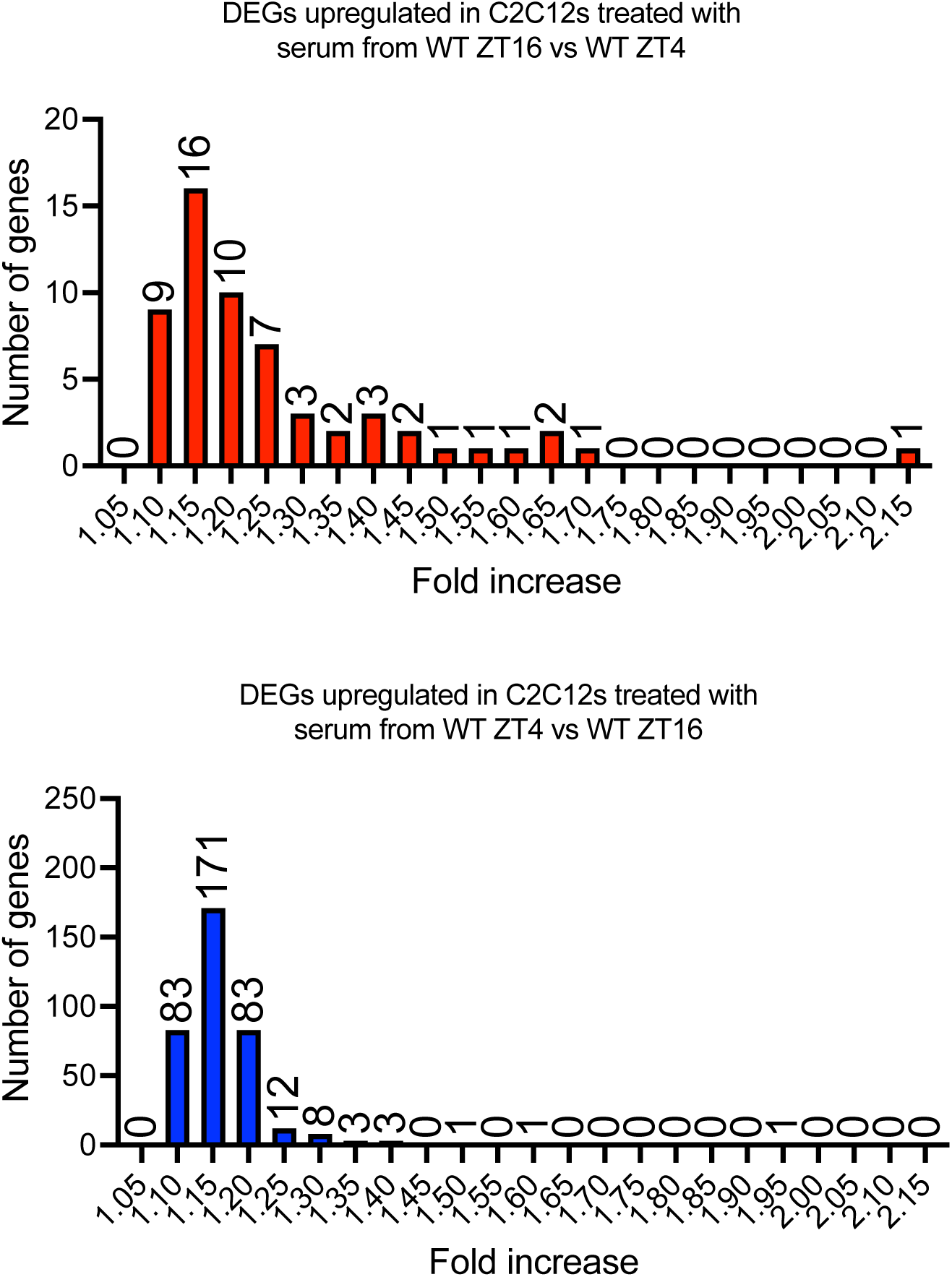
Supporting information for Figure 3. Fold changes of upregulated gene expression (Limma, p<0.01) in C2C12s treated with serum from WT ZT16 vs WT ZT4 (A) and reversed (B).

